# Comparison of dry and wet electrodes for detecting gastrointestinal activity patterns from body surface electrical recordings

**DOI:** 10.1101/2022.04.23.489246

**Authors:** Jonathan C. Erickson, Emily Hassid, Elen Stepanyan

## Abstract

**Objective:** Gastrointestinal motility patterns can be mapped via electrical signals measured non-invasively on the body surface. However, long-term monitoring (≥ 24 hr) may be hindered by skin-irritation inherent with traditional Ag/AgCl pre-gelled (“wet”) electrodes. Therefore, the aim of this work was to investigate the practical utility of using dry electrodes for GI body-surface electrical measurements.

**Approach:** To directly compare dry versus wet electrodes, we simultaneously recorded electrical signals from both types arranged in a 9 electrode (3 × 3) array during an ≈ 2.5 hr colonic meal-response study. Wavelet-based analyses were used to identify the signature post-meal colonic cyclic motor patterns. Signal quality was assessed for each electrode type through quantitative comparison of the dominant frequency, amplitude, signal-to-noise ratio (SNR), and signal energy vs time in the colonic frequency band. Blinded comparison of signal quality was carried out by four expert manual reviewers in order to assess the practical utility of each electrode type for identifying GI activity patterns.

**Main results:** Dry electrodes recorded high-quality GI signals comparable to that of wet electrodes, with dominant frequency in the range 2.85 - 3.25 cpm; peak-peak amplitudes of 120 ± 40 *µ*V, and SNR in the range 7.5 - 11 dB. The CWT colonic frequency band energy versus time correlation coefficient value was ≥ 0.71 for the majority of studies (6 out of 7) indicating very good agreement between dry and wet electrode signals overall. Whereas wet electrodes were rated by expert reviewers as having slightly better signal quality for identifying GI activity patterns, dry electrodes caused no skin irritation and were thus better-tolerated by all subjects.

**Significance:** Dry electrodes are a viable option for long-term GI monitoring studies, offering a potentially more comfortable alternative to conventional wet electrode systems.

## 1. Introduction

Functional gastrointestinal disorders, such as gastroparesis and Irritable Bowel Syndrome (IBS), are extremely prevalent worldwide and associated with substantial health care costs as well as a decreased quality of life [1]. The underlying causes are still under investigation and therapeutic treatment outcomes remain relatively ineffective [2]. Long-term, non-invasive monitoring could substantially aid in the understanding of underlying mechanisms and thus offer more effective targeted treatment options.

The emerging method of non-invasive gastrointestinal (GI) mapping using body surface electrical measurements has recently garnered significant interest for its potential to reveal novel, clinically relevant diagnostic information and therapeutic biomarkers [3]. For example, high-spatial resolution body surface gastric mapping has been used to detect aberrant spatial features of the slow wave correlated to gastroparesis foregut symptoms [4]. A closely related technique, termed electrocolonography (EColG), has been validated to identify cyclic motor patterns in the distal colon evoked by meal ingestion in healthy individuals [5]. EColG could be applied in the future to long-term studies (> 24 hr) of aberrant meal responses in patients suffering from functional disorders such slow-transit constipation [6], as well as monitor the onset and resolution of post-operative ileus over multiple days [7, 8, 9].

GI body surface mapping studies reported in the literature thus far have employed conventional pre-gelled (“wet”) Ag/AgCl adhesive electrodes, which are well-suited for relatively short term recordings (e.g., ≤4 hr meal response studies). However, they may not be particularly well-suited for longer-term recording sessions because the gel dries over time, leading to increased electrode-skin impedance thus lower quality recordings. Furthermore, wet electrodes may be poorly tolerated due to an increased risk of skin irritation and allergic reaction over time [10]. To make long-term GI electrical monitoring studies viable, an alternative electrode platform which overcomes these issues may be required.

Therefore, the purpose of this study was to assess signal quality of dry versus wet body surface electrodes as well as their practical utility for identifying characteristic rhythmic GI activity patterns. Specifically, we investigated whether electrical recordings made with dry electrodes are suitable to identify meal-response colonic cyclic motor patterns (CMPs), in comparison to wet electrodes. Dry electrodes have previously been assessed for their recording quality and overall usability for recording and analysis of the EEG (e.g., [11, 12, 13]), ECG and sEMG [10, 14, 15], but no similar study has yet been conducted for GI electrical signals, which predominantly occur at much lower frequencies, in the range of approximately 0.03 - 0.10 Hz (≈ 2 to 6 cycles per minute (cpm)). The main result of our study was that dry electrodes recorded high-quality signals comparable to those of wet electrodes, and could readily identify colonic CMP post-meal responses in the majority of cases. Thus, dry electrodes may offer a viable alternative electrode platform for long-term GI monitoring applications, in general.

## 2. Materials and Methods

### 2.1. Meal response study

As previously described, colonic CMPs evoked by meal-ingestion can be readily identified using body surface electrical recordings made with wet electrodes [5]. Thus, this hallmark colonic meal response provides a physiologically relevant test bed to assess whether dry electrodes are suitable for measuring GI electrical activity on the skin surface.

We recorded body surface electrical signals generated by the colon during a 2-3 hr meal response experiment, following previous work, with 60–90 min duration pre- and post-meal epochs [5, 16]. In order to minimize pre-meal colonic activity, study participants fasted overnight with oral intake limited only to water after dinner [17, 18]. To transition the colon to an active state, participants ate a meal with size and caloric content matching their daily diet. In healthy normal subjects, meal ingestion induces an rapid and strong increase in cyclic motor patterns (CMPs), which occur predominantly in the rectosigmoid colon, typically lasting for ≥ 30 min immediately after eating, with dominant frequencies in the 2-5 cpm range [16, 19, 5]. The recordings made of colonic signal meal response CMPs allowed direct comparison of dry versus wet electrodes in terms of signal quality and practical utility. All studies were carried out in an academic laboratory setting.

### 2.2. Study participant demographics

A total of 7 recording sessions were analyzed from 3 individuals, 2 of whom participated in 3 separate recordings sessions on different days several weeks apart. One recording from another (fourth) participant was excluded from analysis because identifiable CMPs were recorded by only 2 wet electrodes but none of the dry electrodes, likely owing to poor electrode attachment. All participants self-reported no history of underlying gastrointestinal conditions nor acute GI distress in the 48 hours prior to the recording session. The age (mean ± std) of the study cohort was 20.3 ± 0.6 years old (range = 20 - 21), with a BMI of 22.2 ± 0.8 (range = 21.5 - 23.1)

All participants gave written informed consent, and the studies were approved by the Institutional Review Board of Washington and Lee University.

### 2.3. Electrode arrays

Prior to the start of recording, study participants prepared their abdominal skin surface using Nuprep abrasive skin gel to facilitate low-impedance skin-electrode contact with pre-gelled wet electrodes. It is worth noting that dry electrodes do not generally require skin preparation [15, 12]. Abdominal hair shaving was not performed for any subject.

Study participants self-applied a 3×3 grid of interleaved wet and dry electrodes covering their lower-left abdomen, spaced ≈ 40 mm center-to-center (Figure 1). Verbal instruction from experienced lab personnel was provided, as necessary. The array was positioned to overlay the (recto-)sigmoid colon region, spanning horizontally from the left iliac crest toward the midline, with the upper-most row placed just below the umbilicus. Ground and reference electrodes were placed near the right hip iliac crest.

**Figure 1.**
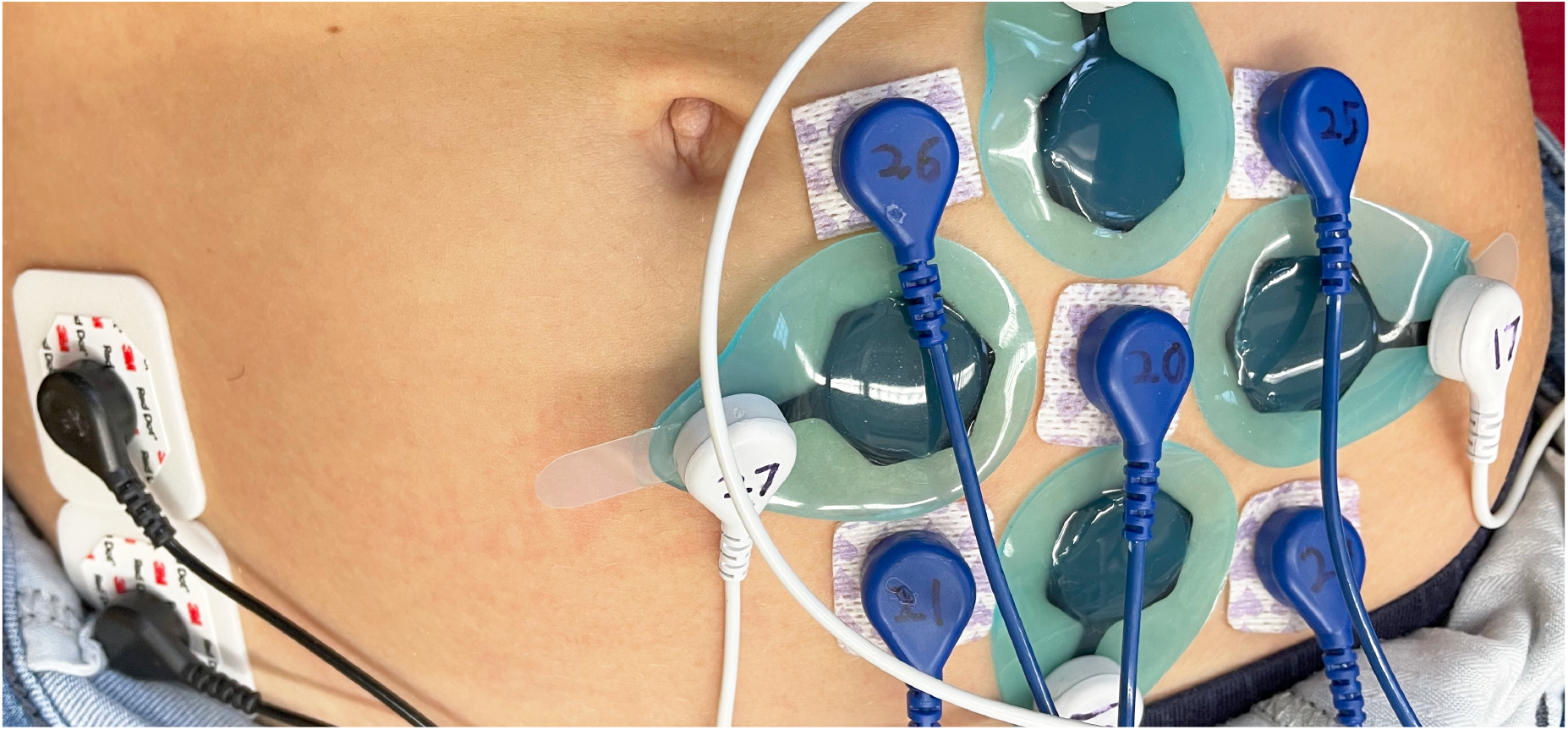
3×3 array for comparing signals recorded with dry versus wet electrodes. The array is positioned approximately above the sigmoid colon. Electrode center-to-center spacing is ≈ 40 mm. Ground and reference electrodes are placed near the right iliac crest. The photo was taken prior to application of surgical adhesive.

Dry electrodes used in this study were soft, skin compliant Dryode Alpha (IDUN Technologies, Zurich, Switzerland; purchased via OpenBCI [20]). They were affixed to the abdominal surface with 3M Tegaderm adhesive for 5 out of 7 recording sessions (labeled numbers 1-4, 7 in Figures 3 and 4; see Discussion). Dryodes incorporate conductive silver microparticles deposited on grasshopper-feet-inspired micropillars which achieve conformal skin contact on micro-and macroscopic scales via van der Waals interactions [15]. The electrical contact area is 20 mm in diameter (≈ 314 mm^2^). The silicone back sheet spans a 55 mm × 40 mm rectangular region (2200 mm^2^). Dryodes in this study were reused for up to 5 meal-response experiments, gently cleaned with 70% ethanol after each use.

Wet electrodes used in this study were conventional pre-gelled with Ag/AgCl conductive adhesive. For 2 studies (recordings numbered 1 and 5 in Figures 3 and 4), we used Danlee Medical brand, model 4500SW, with 546 mm^2^ gel contact area. For the other 5 recordings, we used Ambu BlueSensor P electrodes with 154 mm^2^ electrical contact and 754 mm^2^ total contact area). Wet electrodes were disposed of after use; a fresh set was used for each meal-response experiment.

Wet Ag/AgCl conductive adhesive electrodes were also used for the ground and reference electrodes. During initial recording sessions, these were 3M 2560 red dot (35 mm × 40 mm contact area). However, in latter recording sessions we instead used Ambu BlueSensor P, as some participants noted substantial skin irritation using the 3M brand.

### 2.4. Recording hardware

Electrical signals were recorded using an open-source bioamplifier system [21, 22]. The sampling rate was set to *f*_*s*_ = 100 hz, which was sufficiently high for real-time inspection of the ECG waveform, used here as a proxy measure of good electrode-skin contact. Data files were transferred to a PC for off-line analysis.

### 2.5. Signal preprocessing

Raw recordings for both dry and wet electrodes were analyzed in an identical manner via an automated signal processing pipeline framework that isolates signal components corresponding to colonic CMP activity [5]. Filtering and analysis was performed in the Gastrointestinal Electrical Mapping Suite (GEMS) v2.5 and additional custom software implemented in MATLAB^®^ [23, 24]. Signal preprocessing consisted of 2 main stages: artifact reduction followed by digital filtering as follows.

i. Movement artifact reduction: We adapted and applied the Linear Minimum Mean Square Error (LMMSE) 1-D temporal Wiener filter framework described in a previous work [25] to remove motion and other large transient artifacts. We applied LMMSE using a 30 s data window width to compute statistics averaged over (at least) one colonic oscillation period. Because the signal variance is expected to differ substantially in the post-versus premeal epoch, we used a time-varying noise variance parameter, computed as the running median in sliding 5 min windows, which yields a better time-local noise estimate and thus avoids inappropriate colonic signal attenuation. In brief, this artifact reduction method almost completely removes large brief transients while negligibly distorting the non-artifact segments of the colonic signal component across the full duration of a meal response study.
ii. Band-pass filter: Given the expected dominant frequency of colonic CMPs in the range of 2-4 cpm [16, 8, 5], we implemented a 0.5 - 10 cpm digital bandpass filter (Butterworth, 2nd order, zero-phase). The purpose of the digital filtering step was to remove out-of-band noise and interference such as cardiac and respiration components, as well as other high-frequency noise.

### 2.6. CWT signal analysis

We employed the CWT for spectral time-frequency analysis to quantify changes in rhythmic colonic activity over the duration of the meal response study following previous work [5]. Briefly, the CWT computes a series of complex-valued coefficients **W**(*s, t*) in the time-frequency plane at scales *s* and time-translation *t*. For a discrete data record *x*(*t*_*n*_) with *N* data points the CWT coefficients are computed as:

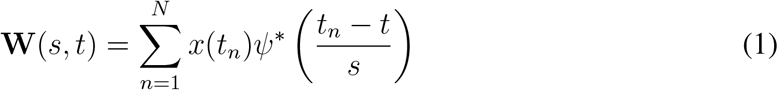

where *ψ* is the normalized wavelet function and ^∗^ denotes the complex conjugate. We used the (approximately) analytic Morlet wavelet 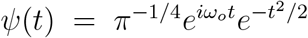 with *ω*_*o*_ = 6. This wavelet was chosen because its shape resembles that of rhythmic colonic components recorded with body surface electrodes. The CWT scales, *s*, can be related to an equivalent frequency 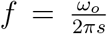, so we may write the coefficients for the electrode indexed by *k* using the notation **W**_*k*_(*f, t*). The CWT was computed using the direct-convolution algorithm with L1-normalization (MATLAB^®^ 2021a function `cwt`), as this yields coefficient magnitudes *W*_*k*_(*f, t*) = |**W**_*k*_(*f, t*)| equal to the amplitude of the underlying signal.

As a global measure of colonic activity for wet and dry electrode subsets, respectively, the mean CWT coefficient magnitude was computed over the *K* electrodes in each subarray within a frequency band centered around the dominant frequency *f*_*dom*_ as follows:

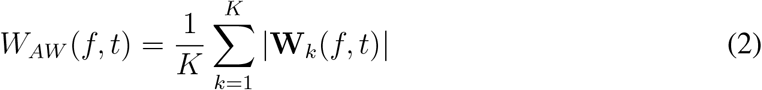

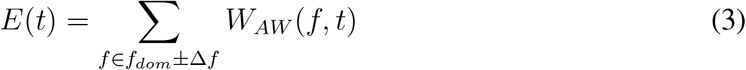

In Eqn. 3, Δ*f* specifies the bandwidth in which CMP signal energy was computed, and the dominant frequency *f*_*dom*_ denotes where the maximal CWT spectral power density, integrated over the full duration experiment duration, occurred:

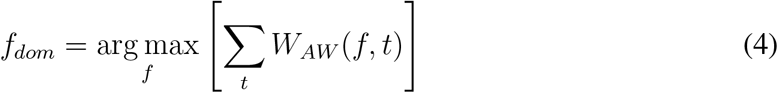

We used a value of Δ*f* = 0.5 cpm based on our observations that a 1 cpm bandwidth is approximately the full-width-half-max of the wavelet frequency spectrum (Figure 2A). Note that *E*(*t*) represents the colonic signal energy as a function of time (Figure 2B).

**Figure 2.**
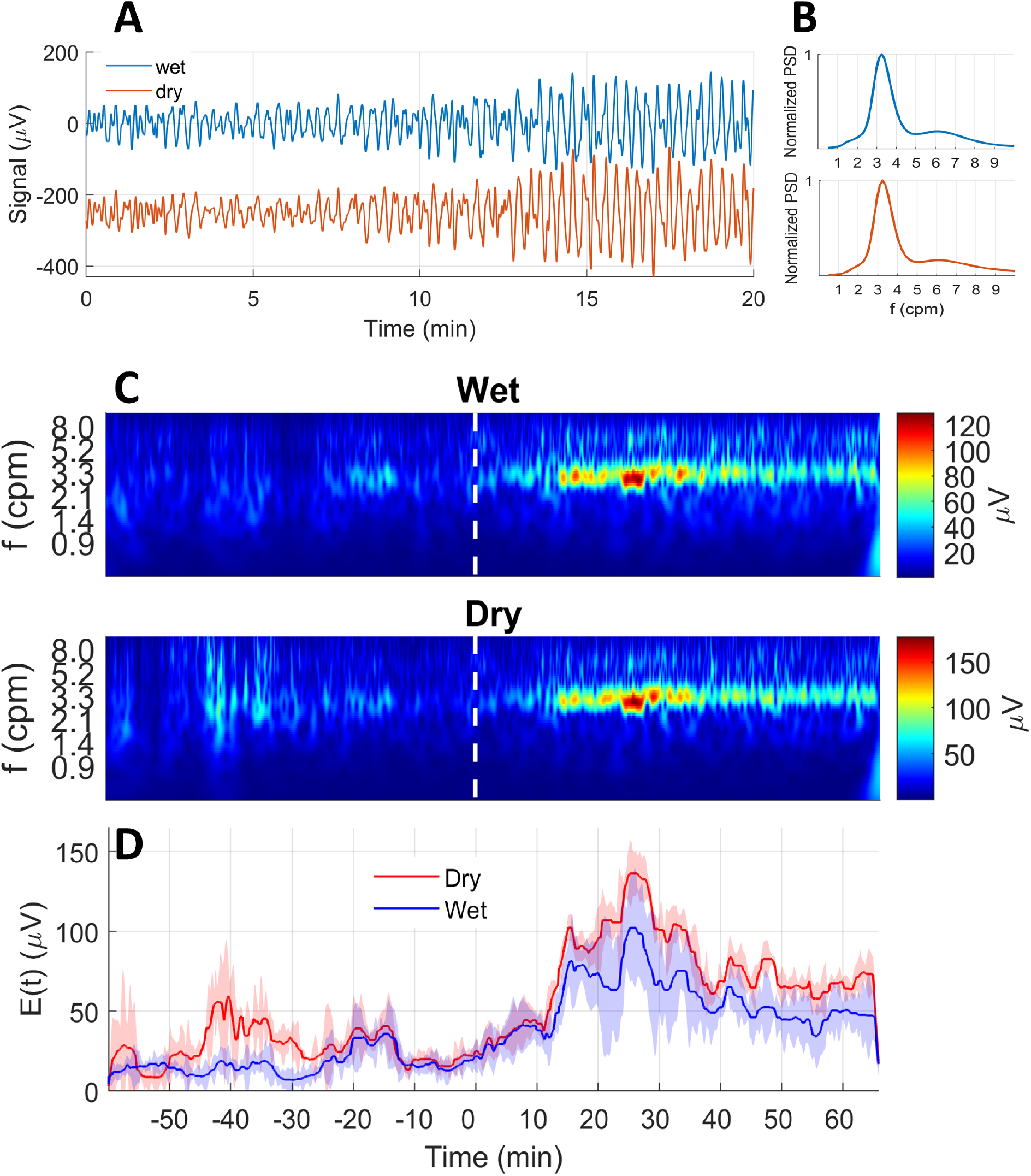
Time-frequency analyses of colonic meal response comparing dry and wet electrodes. Meal ingestion aligned at *t* = 0 sec. A: Sample preprocessed signals for one wet and one dry electrode from a 3 × 3 array. Dry electrode trace is vertically offset for clarity. B: Corresponding power spectral density for wet and dry electrodes are nearly identical, with a dominant frequency 3.24 cpm. C: CWT spectrograms averaged for subsets of wet (top) and dry (bottom) electrodes, respectively. The prominent red horizontal band starting at *t ≈* 15 min indicates a strong meal response recorded by both wet and dry electrodes. Dry electrodes show modestly increased incidence of artifacts during the pre-meal recording (e.g. *t ≈* −40*to* – 30 min). D: Amplitude of colonic component vs time showing comparable time course for meal response. Dry electrodes measured a larger amplitude during peak post-meal response in this recording.

As the goal of analyses was to quantify identifiable changes in the colonic signal component in the fasted versus fed states, two fundamental quantities of interest were the pre-meal baseline signal level *E*_*pre*_, and the signal amplitude during periods of colonic activity *E*_*active*_. These were quantified for dry and wet electrode subsets, respectively, as follows:

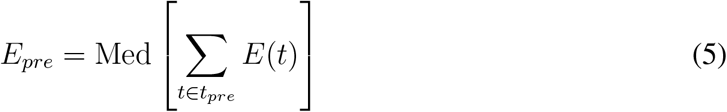

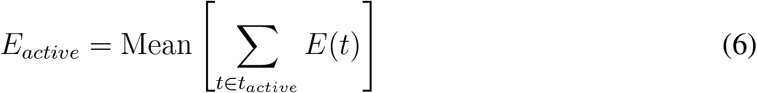

In Eqn. 5, the time subscript *t* ∈ *t*_*pre*_ denotes the set of time points in the pre-meal epoch. In Eqn. 6, *t*_*active*_ denotes the set of time points in the post-meal epoch at which the colon was deemed “active”. These were determined in automated fashion by finding all time-points in the post-meal epoch for which *E*(*t*) > *E*_*pre*_ ± 2.5 *σ*_*pre*_, where *σ*_*pre*_ represents the statistical variance (noise) in the pre-meal colonic signal, estimated using the median of the absolute deviation. In practical terms, the factor of noise multiplier of 2.5 chosen here requires that active time points exhibit colonic frequency band energy ≥99% percentile value observed during the pre-meal epoch.

We used the median for computing *E*_*pre*_ to obtain a robust estimate of the “true” baseline colonic signal energy, not strongly biased by the presence of motion artifacts. Note that doing so excludes any colonic signal which may be attributed to a pre-meal cephalic response. The mean was used to robustly estimate the colonic signal amplitude in the post-meal epoch, given that the time points of colonic activity were already explicitly identified. Motion artifact components which may not completely reduced during preprocessing may still contribute to this computation, but their overall effect is expected to be minor because they are short-lived, and their amplitude is no greater than that of the colonic CMPs.

### 2.7. Array-wide CWT signal quality comparison

To quantitatively compare the signal quality of CMPs recorded with each electrode type, we computed the Pearson correlation between array-wide CWT spectrogram coefficient magnitudes *W*_*AW*_ (*f, t*) (Eqn 3). This essentially computes a similarity score between corresponding pairs of wet and dry electrode CWT spectra, for example those shown in Figure 2C.

Additionally, we computed correlation coefficients for each corresponding wet-dry pair of colonic frequency band signal vs time *E*(*t*) (Eqn 3). This measures the degree to which CMPs may be readily identified from the brightly colored horizontal bands within a restricted frequency range, for example the time series for wet and dry electrodes illustrated in Figure 2D.

### 2.8. Individual electrode signal quality and usability comparison

To further assess individual signal quality and compare the practical utility between electrode types, we computed the following physiologically relevant metrics for each individual electrode’s CWT spectrogram:

i. Dominant frequency (*f*_*dom*_). These were similarly determined per Eqn 4, but using CWT magnitude spectra for individual electrode signals (e.g. Figure 2A) instead of array-wide averaged spectrograms.
ii. Peak-to-peak colonic signal amplitude (*E*_*pp*_):

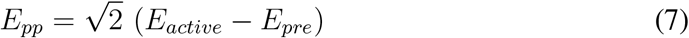 This estimate subtracts the baseline signal level *E*_*pre*_ from the active signal amplitude *E*_*active*_ to obtain the pure colonic component only. The 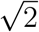 factored into *E*_*pp*_ accounts for the fact that *E*_*active*_ and *E*_*pre*_ are essentially RMS estimates as they are derived from summing across wavelet coefficients in the colonic frequency band which fall off monotonically on either side of the dominant frequency.
iii. Percent change in colonic CMP signal amplitude in the fed state relative to fasted (Δ*E*):

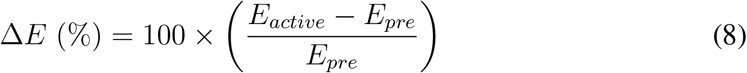
iv. Signal to noise ratio (*SNR*):

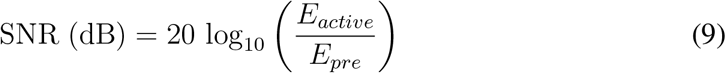 This estimate assumes that the colon is quiescent during the majority of the pre-prandial epoch; thus *E*_*pre*_ may be considered as noise only.

### 2.9. Qualitative assessment: signal quality and interpretation

Four expert reviewers inspected in a blinded manner paired wet versus dry CWT spectra for each of 7 recordings (e.g., Figure 2C) and were asked to rate the following: 1) overall signal quality of the CWT spectrograms for each of 7 subjects; and 2) similarity of underlying colonic CMP activity identified from visual interpretation of the spectrograms. The latter serves as an overall indicator of practical utility.

Both signal quality and visual interpretation were rated on a 0-3 magnitude scale with discrete scores corresponding to differences of: 0 = same/no difference; 1 = slight/minor; 2 = modest; 3 = large and substantial. A positive score indicated the dry electrodes offered superior quality recordings; a negative score indicated wet electrodes were rated higher.

### 2.10. End-user comfort

Each study participant was asked to rate using a 7-point scale their experience comparing wet and dry electrodes in terms of ease of application and overall comfort. A higher score indicated better performance. For ease of application, a score of 1 corresponded to “Difficult and/or time consuming” while a 7 indicated self-application of the electrode was “As easy as applying a Band-Aid.” For overall comfort, a score of 1 corresponded to “Uncomfortable, substantial deterrent to prolonged use” whereas 7 indicated“Comfortable/no bother”. Additionally, participants were also invited to provide additional comments that addressed these points.

### 2.11. Statistical methods

Quantitative metrics are reported below as the mean ± s.d., unless otherwise specified. The Mann-Whitney U test was used for statistical comparison for wet versus dry electrode metrics described in Section 2.8 This compares the median value of the distributions, with a significance threshold based on a 95% confidence level; *p* ≥ 0.05 indicated median values were statistically indistinguishable.

## 3. Results

Dry electrode signal quality was generally very good and comparable to that measured with wet electrodes. There were 2 instances in which statistically significant differences noted, which are also easily identified from visual analysis. These include the peak-to-peak signal amplitude *E*_*pp*_ for recording 6 and the percent change in energy Δ*E* for recording 7.

### 3.1. Quantitative metrics

Figure 3 summarizes the panel of metrics comparing dry versus wet signal quality obtained from individual electrode analyses (panels A-D) and using array-wide CWT spectral analysis (panels E and F).

**Figure 3.**
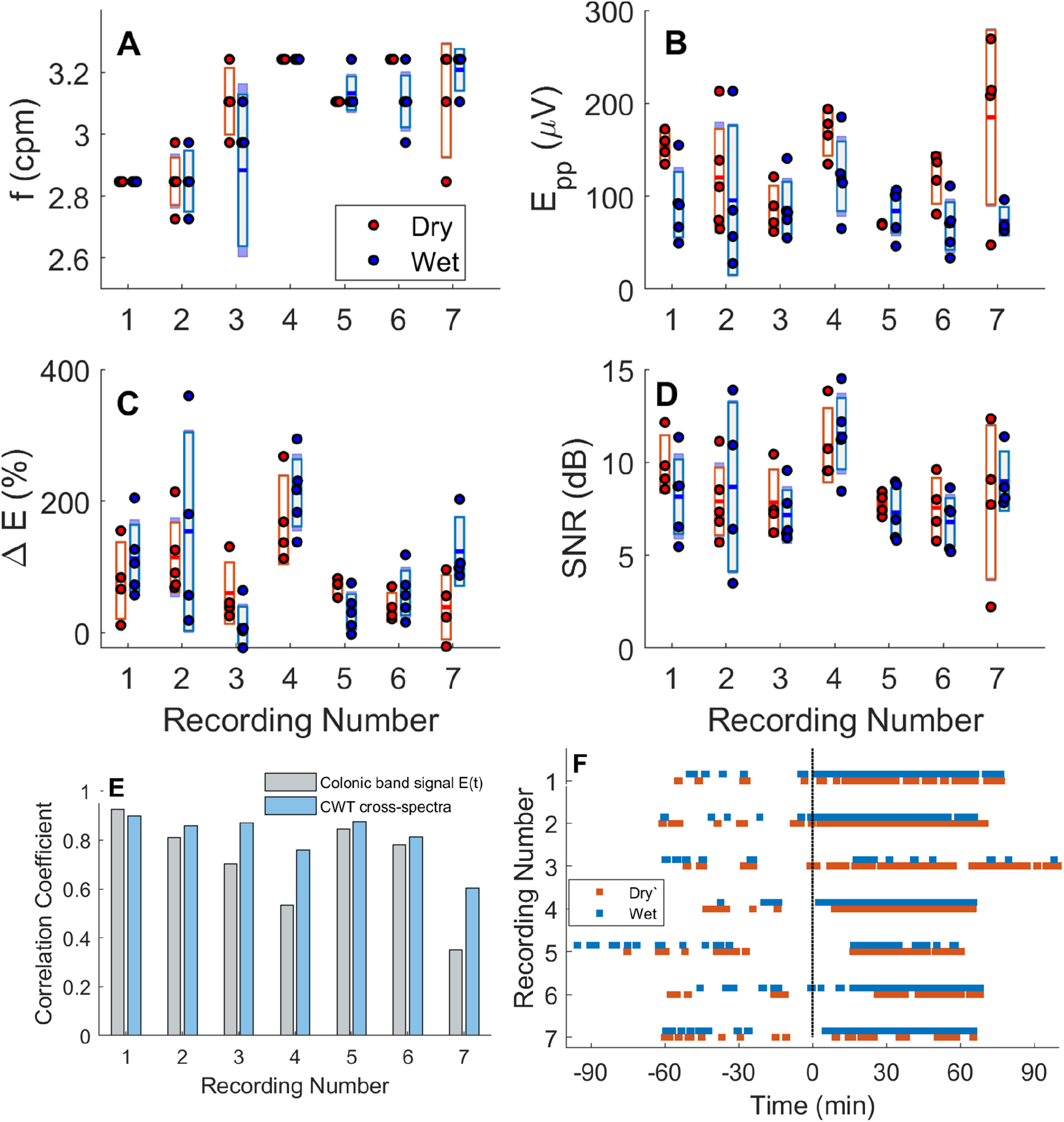
Box plots for various metrics. Individual electrode responses are plotted as dots. The metrics are dominant frequency (A); Peak-to-peak amplitude (B); Amplitude change in post- vs pre-meal epoch (C); Signal-to-noise ratio (D). Cross-CWT spectra and colonic frequency band signal vs time correlation coefficients (E); automated CMP activity lines indicate similar outcomes for some but not all recordings (F).

i. Dominant frequency (Figure 3A) showed very good agreement between dry and wet electrodes. Both dry and wet electrode arrays yielded dominant frequencies of 2.85 - 3.25 cpm, falling within the canonical colonic CMP frequency range [16, 19]. The maximum difference observed in the mean and median dominant frequencies between the two electrode types was 0.11 cpm (recording 6), not deemed in this case to be of particular physiological importance.
ii. Peak-to-peak CMP signal amplitude (Figure 3B) also indicated robust signal quality for dry and wet electrodes. Recording-specific median *E*_*pp*_ values of 119 ± 40.1 (range= 69 - 153) *µ*V were recorded with dry electrodes—excluding recording 7, whose dry electrode recordings were observed to be contaminated by motion artifacts (see discussion)—compared to 85.4 ± 18.5 (range = 66 - 118) *µ*V for wet. The difference in amplitude observed might be due to higher impedance values associated with wet electrodes compared to Dryodes at low frequencies [15] of ≈ 0.05 Hz (3 cpm) leading to a higher pre-meal baseline noise level and thus decreased estimates for colonic signal amplitude.
iii. The expected large postmeal increases in the colonic band signal amplitude were observed for both dry and wet electrodes across all recordings (Figure 3C). The recording-specific median Δ*E* was 72.3 ± 41.6% (range = 32.3 - 153 %) rendering colonic CMP meal responses easy to identify by computer-automated algorithms and human visual analysis for the majority of recordings. Wet electrode signal quality achieved higher values of 90.9 ± 69.7 %, (range = 30.6 - 218 %), indicative of better signal quality.
iv. The signal-to-noise ratio for dry and wet electrode SNR were comparable for all 7 recordings (Figure 3D). The observed recording-specific median SNR for dry electrodes ranged from 7.5 - 10.9 dB versus 7.2 - 11.5 dB for wet electrodes, with an intra-recording average difference of only 0.9 ± 0.4 dB.
v. CWT-based correlation coefficients (Figure 3E) indicate a high degree of similarity for 5 out of 7 recordings; moderate agreement for 1 recording; and poor agreement for the last one due to relatively poor dry electrode performance. The colonic frequency band *E*(*t*) median correlation coefficient value was 0.71 ± 0.20 (median = 0.78; range = 0.46 - 0.94). The CWT cross-spectrum correlation measured across all 7 recordings was of 0.81 ± 0.10 (median = 0.86; range = 0.64 - 0.91). Taken together, these indicate concordant results obtained with both dry and wet electrode types for the majority of recordings. For one case (recording 4) where a lower correlation coefficient values was observed, the signal quality and amplitude was sufficiently high for both dry and wet electrodes such that there were negligible differences in the overall CMP activity pattern identified from recordings of either type. However, for another (recording 7), some electrodes in the dry electrode array failed to record high quality signals, likely due to the fact the subject presented with a hirsute abdomen leading to poor electrode-skin contact.
vi. CMP activity lines across all 7 recordings (Figure 3F) indicated 78 ± 9.8 % agreement overall (range = 61 - 84 %) between wet and dry electrodes. Again, for 5 recordings, the post versus pre meal CMP activity indicated is very similar, while for the other 2 recordings there are noticeable differences. For recording 3, the dry electrodes appear to more reliably track the expected progression of post-meal colonic CMPs, whereas the wet electrodes appear to do so for recording 7.

### 3.2. Qualitative signal and practical utility assessment

Figure 4 summarizes the expert reviewer scoring regarding the signal quality and similarity of colonic motor patterns visually identified using dry electrode signal analysis compared to wet. The latter may be regarded as a measure of practical utility.

**Figure 4.**
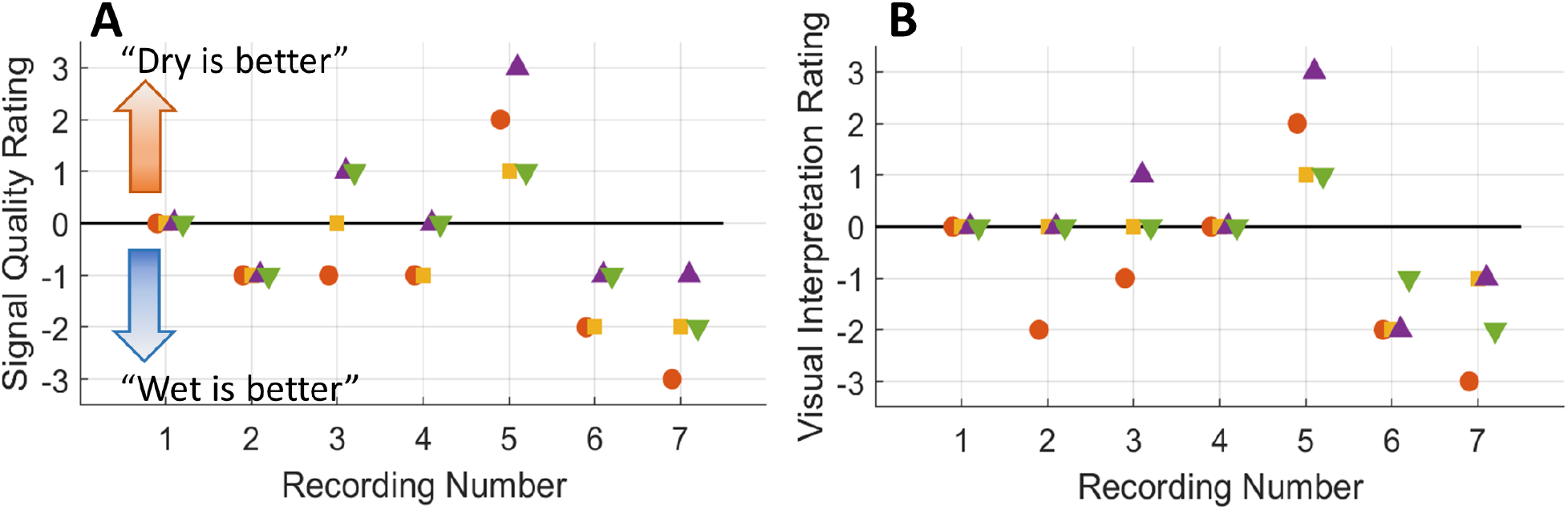
Human reviewer ratings for CWT spectrogram quality (A) and similarity of colonic CMP activity identified visually from CWT spectrograms (B). Wet electrodes were rated as having modestly better performance overall.

Dry electrodes were rated as having similar though slightly inferior signal quality overall, with a rating across all 7 recordings of −0.48 ± 1.4 (on a 3 point scale), with a median = -1. For 2 recordings (numbers 1 and 3), the signal quality was rated essentially the same (median score of 0). For 2 participants (recordings 6 and 7), the wet electrodes were rated as having substantially better signal quality compared to dry (median score of 2). The ratings for the 7 recordings indicated only slight differences (median score of 0), with 1 case where dry electrodes offered better signal quality in 1 instance (recording 5) compared to 2 instances for wet electrodes (recordings 2 and 4).

In terms of practical utility, dry electrodes were also rated as modestly inferior for visual identification and interpretation of CMP activity. The score across all 7 recordings was −0.33 ± 1.4 (median = -1.4). There were 3 cases (recordings 1, 3, and 4) where no overall difference was observed (median score of 0); 2 participants (recordings 2 and 7) where a modest discrepancy was noted (median score of 1); and 2 cases (recordings 6 and 7) where substantial differences were noted.

Overall, these results indicate while wet electrodes provided overall better signal quality and clearer visual interpretation of CMPs, dry electrodes are suitable, in a majority of cases, for identifying rhythmic GI motility patterns from body surface electrical recordings.

### 3.3. End-user ease of application and comfort

Participants uniformly reported that whereas wet electrodes were easier to apply, dry electrodes were more comfortable and better tolerated overall.

The ease of application score was higher for wet electrodes (6.0 ± 1.0) compared to Dryodes (3.3 ± 0.58). By contrast, the comfort ratings evidenced a clear preference for dry electrodes (6.33 ± 0.58) over wet (4.0 ± 1.0). Users reported dry electrodes required more dexterous and time-consuming manipulation to achieve good contact.

Wet electrodes used in this study produced dermal irritation over the duration of the meal study, with 2 study participants describing uncomfortable and noticeably inflamed skin upon electrode removal. By contrast, dry electrodes produced no skin irritation, and were comfortable and well-tolerated over the full duration of the meal study. This is a key difference between dry and wet electrode types.

## 4. Discussion

To our knowledge, this study is the first to asses whether dry electrodes are suitable for GI body surface electrical measurement, in comparison to wet electrodes. The results show that dry electrodes can achieve excellent signal quality that is sufficient for identifying colonic post-meal CMPs in a controlled laboratory environment. Although dry electrodes achieved sufficiently high-quality recordings to accurately identify colonic CMPs in a majority of cases (6 out of 7), it is worth noting that expert human reviewers rated wet electrode as recording modestly superior quality signals overall.

Whereas wet electrode signal quality was observed to slightly degrade over the course of a 2-3 hr experiment, dry electrode signal quality was often observed to improve over a time frame of 10-30 minutes and remain stable thereafter, owing the adhesion mechanism yielding decreased skin-electrode impedance over time [14, 15]. Dry electrodes appear to be advantageous for longer-term monitoring of GI electrical activity patterns in this regard.

In this study, participants comfortably rested in a supine position for the duration of the experiment in order to facilitate good skin-electrode contact and to minimize the occurrence of motion artifacts. Even so, dry electrode recordings were observed to be more prone to motion artifacts, likely due their less robust method of attachment. While computer-automated algorithms can reduce artifacts in GI electrical recordings (e.g. [25, 26]), a future study should address how to further minimize their impact, which may otherwise be problematic for general application in an ambulatory setting.

A method to maintain long-term mechanical attachment of dry electrodes to facilitate stable, low-impedance electrical contact must be established in order to make long-term studies feasible—preferably without the use of a surgical adhesive. In this study, Tegaderm was not used to affix dry electrodes for 2 recordings (labeled numbers 5 and 6 in Figures 3 and 4) to minimize the risk of dermal discomfort in a participant who reported that it caused mild skin irritation.

Additionally, Dryodes commercially available at the present time would need to manufactured with a reduced footprint to be compatible with high spatial density mapping platforms used to map propagating waves associated with GI motility, i.e., the gastric slow wave [27, 28]. One potential method for surmounting these current challenges would be to integrate Dryode technology into stretchable, conformal soft electronics platforms for GI application [29, 30], or into elastic textile garments [31].

A limitation of the current study was the demographic range of the study participants. (It is worth noting that study participant recruiting was restricted per COVID-19 safety protocols in place at the time this study was performed.) It is plausible to hypothesize that the results obtained in this study would generalize to a larger BMI values ≤ 27 on the basis that colonic CMP meal responses were previously identified in this range [5, 32].

In this study, we made comparisons using several types of wet electrodes with varying contact area, and the wet and dry electrode contact areas also differed by a factor of ≈ 2×. We did not observe systematic differences in results obtained using different contact area wet electrode types, nor would we expect to observe such differences based on biophysical principles. First, given the diameter of the distal colon—the underlying electrical source—has a diameter of ≈ 50 mm subject to the volume-conduction spatial low-pass spatial filtering, the body surface potential would spatially vary by only ≤ 5% over the contact area of any electrode employed in this study [33]. Second, although we did not directly measure the skin-electrode impedance, it is reasonable to assume values were ≤ 1MΩ at a frequency of ≈0.05 Hz for all electrode sizes [15]. This is 100× less than the 100 MΩ amplifier input impedance. Therefore, the voltage divider effect would be negligible.

Finally, the current study used only wet electrodes for ground and reference points. An ideal study design would implement two separate amplifier systems operating simultaneously, each with dedicated wet and dry grounding configurations, respectively. A previous study assessing dry electrode EEG sleep recordings referenced to either wet or dry electrodes revealed only minor differences in signal quality [12]. Thus, we would not expect to observe significant differences dependent upon on the reference electrode type when measuring GI electrical signals.

## 5. Conclusion

This study offers the first comprehensive comparison of dry versus conventional pre-gelled wet electrodes for cutaneous measurement of gastrointestinal electrical signals. The results show that Dryodes are suitable for clearly identifying colonic post-meal response CMPs. Signal quality obtained with dry electrodes was very good, similar to that of wet electrodes, and characteristic post-meal colonic CMPs easily identified in the majority of recordings. While users found dry electrodes somewhat more time consuming and difficult to self-apply, they were also rated as more comfortable and tolerable for long-term studies compared to wet electrodes. The use of dry electrodes may be extended to include measuring gastric slow wave activity on the body surface, given the similarity in colonic and gastric amplitude and dominant frequencies. Dry electrodes may thus facilitate non-invasive, long-term monitoring of GI electrical activity to further elucidate dynamic motility patterns in health and disease.

## 6. Acknowledgments

We thank the expert manual reviewers for offering their candid assessment and interpretation of signal quality and study outcomes.

